# MeQTL Mapping in African American Hepatocytes Reveals Shared Genetic Regulators of DNA Methylation and Gene Expression

**DOI:** 10.1101/2025.01.23.634506

**Authors:** Kathryn Carver, Carolina Clark, Yizhen Zhong, Guang Yang, Mrinal Mishra, Cristina Alarcon, Minoli Perera

## Abstract

Methylation quantitative trait loci (meQTL) mapping can provide insight into the genetic architecture underlying the epigenome by identifying single-nucleotide polymorphisms (SNPs) associated with differential methylation at methylation sites (CpGs) across the genome. Given that the epigenetic architecture underlying differences in gene expression can vary across racial populations, performing epigenomic studies in African Americans is crucial for understanding the interplay between genetic variation, DNA methylation, and gene expression in this understudied group. By performing cis-meQTL mapping in African American hepatocytes, we identified 410,186 cis-meQTLs associated with methylation at 24,425 CpGs in the liver. Through colocalization analysis, we found that 18,206 of these meQTLs are also colocalized with known liver eQTLs. Additionally, we found that using African American eQTL data results in an increased ability to detect additional colocalized variants that exhibit strong differences in allele frequency between people of European and African ancestry. Furthermore, the presence of smaller linkage disequilibrium blocks in African Americans allows us to identify narrower genomic regions of potentially causal variants compared to when data from Europeans is used. Importantly, these colocalized SNPs are significantly enriched for genetic associations with lipid and inflammatory traits in the GWAS catalog, suggesting that DNA methylation may contribute to the etiologies of these diseases. Furthermore, while it is generally presumed that the genetic regulation of DNA methylation is shared between blood and liver, we found that only 5.4% of African American liver meQTLs colocalize with blood meQTLs. Overall, our results reveal that studying African American populations results in the identification of additional genetic and epigenetic factors that may regulate gene expression in the liver, thereby expanding our understanding of gene regulation in African Americans.

## Introduction

Genome-wide association studies (GWAS) have become a powerful tool for understanding the role of genetic variation in regulating disease pathogenesis, drug response, and other clinically relevant traits. Nearly 90% of variants previously identified in these studies lie in non-coding regions of the genome, making their role in disease pathogenesis unclear.^1^ While these variants do not directly alter the structure of the protein encoded by that gene, they can have significant effects on disease and drug response by regulating gene expression through epigenetic mechanisms.^2^ DNA methylation, an epigenetic modification to DNA that is typically associated with decreased gene expression, has been implicated in a broad range of diseases, including autoimmune disorders, metabolic disorders, hematological malignancies, and cancer.^3^ DNA methylation near genes involved in drug metabolism has been found to influence pharmacokinetics and pharmacodynamics.^4^ Given that the liver is a critical organ for drug metabolism and is involved in diseases such as metabolic conditions, cardiovascular disease, and cancer, identifying genetic and epigenetic regulators of liver gene expression is crucial for understanding these phenotypes.^5, 6, 7, 8^

Identifying genetic variants associated with differences in DNA methylation through methylation quantitative trait loci (meQTL) mapping can reveal the genetic architecture underlying individual differences in the epigenome.^9^ Furthermore, integrating meQTL mapping with other multi-omic analyses, such as expression quantitative trait locus (eQTL) mapping, can clarify how interactions between genetic variation, epigenetic variation, and gene expression contribute to disease pathogenesis.^9^ However, most meQTL and eQTL studies performed so far have focused on European populations.^10^ Minority racial populations, especially African Americans, are severely underrepresented in this field of research. Given that African populations possess additional genetic variation not found in Europeans, the causal variants regulating gene expression in this population may be unique from the causal variants identified in other populations.^11, 12, 13^ Additionally, the epigenetic architecture underlying gene expression can vary across populations.^9^ Thus, performing genomic and epigenomic studies in African Americans, particularly in the liver, is critical for elucidating the mechanisms regulating gene expression, disease risk, and drug response in this population.

Here, we present the largest meQTL mapping study performed in African American hepatocytes to date. To investigate the role of these meQTLs in regulating gene expression, we used colocalization analysis to integrate this data with eQTL data from the GTEx liver dataset and an eQTL dataset from American American hepatocytes. To better understand the role of these SNPs in disease pathogenesis, we analyzed their direction of effect, compared our results to previously identified African American blood meQTLs, and calculated enrichment for GWAS EFO categories. Together, these results represent a significant step in characterizing the genetic and epigenetic architecture underlying gene expression in African American hepatocytes.

## Methods

### 1. Cohort Description

A total of 79 African American primary hepatocyte cultures were used for this study. Hepatocytes were obtained from companies (BioIVT, TRL, Life Technologies, Corning, Xenotech) or isolated from cadaveric livers using the procedure previously described in Park et al.^14^ Livers with active cancer or a history of hepatocarcinoma were excluded from analysis. 77 samples were used for meQTL analysis due to the availability of paired DNA methylation and genomic data. Of the 77 total samples used, 38 were derived from female donors and 39 were derived from males. Donor ages ranged from 18 years to 72 years old with the average age being 45.8 years.

### 2. Genotyping and Imputation

DNA was extracted from hepatocytes as described in Park et al.^14^ SNPs were genotyped using the Illumina Multi-Ethnic Genotyping Array. We used standard pre and post-imputation quality control measures as described in Zhong et al.^10^ After quality control and imputation, 8,442,281 SNPs remained in the study. To identify and correct for hidden variables in the genotype data, we performed Principal Component Analysis (PCA) as described in Zhong et al.^10^ Two genotype PCs, PC1 and PC2, and age and sex were included as covariates in downstream analyses.

### 3. Methylation Profiling and Quality Control

DNA was isolated from hepatocytes as described in Park et al.^14^ 79 samples produced sufficient bisulfite-converted DNA for methylation analysis. As described in Singh et al, the Illumina MethylationEPIC BeadChip microarray was used to profile methylation levels at approximately 850,000 methylation sites across the genome.^15^

Methylation quality control and normalization procedures were performed using the *ChAMP* R package.^16, 17^ Quality control removed 10,798 probes for any sample that did not have a detection p-value <0.01, 1,832 probes with a bead count <3 in at least 5% of samples, 2,968 probes with no CG start sites, 95,322 probes with SNPs as identified in Zhou et al.^18^, 11 probes that align to multiple locations as identified in Nordlund et al,^19^ and 16,474 probes located on X and Y chromosomes. To fix outliers in the dataset, all values smaller than or equal to 0 were replaced with the smallest positive value while all values greater than or equal to 1 were replaced with the largest value below 1. After QC, 738,513 probes and all samples remained in the study. Following probe QC, the methylation data was normalized and corrected for batch effects. To adjust for type-II probe bias, we performed type-II probe normalization using the BMIQ method.^20^ Next, we corrected for batch effects using the singular value decomposition method implemented by Teschendorff et al. to identify significant components of variation in the methylation data.^20^ Ideally, variation in a methylation dataset can be attributed to biological factors. However, technical sources of variation can introduce additional variation into the dataset. ChAMP was used to identify technical factors correlated with variation in the dataset and reduce the effects of confounding variables, such as which slide or batch each sample was analyzed in (Supplementary Figure 1). We used the ComBat function to correct for these technical variables as well as biological variables such as age and sex. Of the 79 samples that underwent methylation profiling, 77 also had genotype data. These samples were used for downstream analyses.

### 4. Methylation Quantitative Trait Loci Mapping

We performed methylation quantitative trait loci (meQTL) analysis for 77 samples that passed both genotype and methylation quality control procedures. We performed meQTL analysis using sex, age, and two genomic PCs as covariates. A total of 8,442,281 SNPs and 738,513 methylation sites were analyzed. We performed cis-meQTL analysis using a cis window of 50 kB in *MatrixEQTL* in R.^21^ To call significant meQTLs, we used the hierarchical correction method described in Zhong et al with a false discovery rate threshold of 0.05.^10^

### 5. Colocalization Analysis

Following meQTL analysis, we performed colocalization analysis using the *coloc* package in R.^22^ To identify SNPs coregulating methylation and gene expression in the liver, we colocalized significant liver meQTLs with two liver eQTL datasets (GTEx Liver V8 and African American eQTLs published in Zhong et al).^23, 10^ Because the same SNP can be associated to the expression of multiple genes, we iteratively performed colocalization analysis for each gene separately. For a given gene in the eQTL dataset, we identified all SNPs that were eQTLs for that gene. From the meQTL data, we then pulled all meQTL associations for these SNPs. We then performed colocalization between the overlapping significant meQTL and eQTL SNPs for each eGene. This process was repeated for every eGene in each eQTL dataset. We defined significant colocalizations as those having a PP4 > 0.80. We also performed colocalization analysis using the local ancestry-adjusted eQTL dataset from Zhong et al., which did not identify a significant number of additional colocalized signals.^10^

To identify SNPs regulating methylation in both liver tissue and blood, we colocalized significant liver meQTLs with significant blood meQTLs from the GENOA study.^9^ For a given significant CpG, we performed colocalization between SNPs that were associated with this CpG in the liver dataset and blood dataset. We repeated this process for every significant meCpG. As previously mentioned, PP4 > 0.80 was defined as the threshold for colocalization. We did not perform any pruning before performing colocalization with either dataset. If a SNP was associated with multiple CpGs and exhibited colocalization with multiple CpGs, we only reported the lead association. The DAVID NIH tool was used to perform gene enrichment and KEGG pathway analysis following colocalization.^24^

### 6. Mediation Analysis

Following colocalization, we used the *mediation* R Package to perform mediation analysis and determine whether methylation mediates the effects of genotype on gene expression for colocalized meCpG-eGene pairs.^25^ We performed this analysis using 53 individuals who have genotype, methylation, and gene expression data. Mediation analysis was conducted by testing the effect of methylation in two directions. First, we performed SNP-Methylation-Expression (SME) analysis where SNP genotype is the exposure, methylation is the mediator, and gene expression is tested as the outcome. Second, we performed SNP-Expression-Methylation (SEM) analysis where SNP genotype is the exposure, gene expression is the mediator, and methylation is tested as the outcome. For both types of mediation analysis, we adjusted for age, sex, the top 2 genomic PCs, and the top 4 expression PCs. Using the *mediation* package in R, we used the same regression formulas as described in Shang et al. to test the SME and SEM models^9^. We performed bootstrapping with 1000 simulations to calculate the mediation proportion and p-value. We declared significance at a FDR of 0.05. <u>SNP/CpG Annotations and Enrichment Analysis</u>

Following meQTL mapping and colocalization analysis, we annotated SNPs to their respective genomic regions (5’ UTR, promoter, gene body, etc). We performed these annotations using ANNOVAR in the Functional Mapping and Annotation of Genome-wide Association Studies (FUMA-GWAS) software.^26^ We calculated enrichment statistics for significant meQTLs and colocalized SNPs by performing Fisher’s Exact Test against the 1000 Genomes ASW reference panel used to calculate SNP enrichment statistics in the FUMA-GWAS documentation.^26^ To obtain annotations for CpG sites, we used the annotation functions embedded in the *ChAMP* R package.^16^

#### GWAS Enrichment Analysis

Following colocalization analysis, we also calculated GWAS EFO term enrichment for meQTLs colocalized with the African American eQTL dataset. We used all statistically significant SNP associations in the NHGRI-EBI GWAS catalog^27^ for this analysis. As done in Zhong et al., we extracted all variants in LD with the independent GWAS variants and colocalized variants (r^2^ > 0.8) using African Americans as the LD population.^10^ We chose this approach because there are known LD differences between our African American meQTL and eQTL data compared to the GWAS catalog data, which is primarily composed of individuals of European ancestry.

To determine if our colocalized SNPs were enriched for particular GWAS parent EFO categories, we randomly sampled a null set of 1000 SNPs matched with our meQTLs for MAF, distance from the nearest TSS, and LD score within 10 quantile bins, as done in Zhong et al.^10^ To test for enrichment, we identified the number of null SNPs overlapped with a significant association in the GWAS catalog and averaged the number of overlapped SNPs falling into each parent EFO category across 1000 random sets. We then identified colocalized SNPs that are overlapped with a significant association in the GWAS catalog, counted the number of SNPs that fell under each parent term, and compared that to the average number of null set SNPs falling in that parent category using Fisher’s Exact Test. We corrected the p-value for Fisher’s Exact test using the Benjamini-Hochberg (BH) procedure and set a threshold for significance at an adjusted p-value of 0.05.

## Results

### MeQTL Analysis

To identify SNPs associated with differences in methylation at CpG sites across the genome, we performed methylation quantitative trait loci (meQTL) mapping. Because we were most interested in examining the effects of SNP variation on local methylation, we performed cis-meQTL analysis using a cis window of 50 kB. MeQTL analysis identified a total of 410,186 meQTLs, with these SNPs being associated with differential methylation at 24,425 unique methylation sites (CpGs).

Of the 24,425 CpGs identified, 14,423 (59.1%) fell into open sea regions (Figure 1A). 4,620 (18.9%) CpGs were found in the promoter near transcriptional start sites (1,296 within 200 bp and 3,324 within 1500 bp of the TSS), indicating a potential regulatory role in gene expression (Figure 1B). For the 410,186 SNPs found to be associated with CpG methylation, we observed a statistically significant enrichment for SNPs located in the 5’ UTR, 3’ UTR, intronic regions, upstream regions, ncRNA exonic regions, downstream regions, and exonic regions (Figure 1C). We observed a statistically significant depletion of SNPs in ncRNA intronic and intergenic regions (Figure 1C). Most SNPs associated with differences in methylation are near the CpG they affect with 27.9% of meQTL SNPs within 5 kB of the CpG to which they are associated and 43.7% within 10 kB (Figure 1D). There is a high density of meQTL SNPs located proximally to the CpG sites they are associated with, while there are relatively few SNPs that are significantly associated with CpG sites greater than 25 kB away. SNPs associated with increased methylation (positive beta) exhibit the same distribution as SNPs that are associated with decreased methylation (negative beta) (Figure 1D).

**Figure 1:**
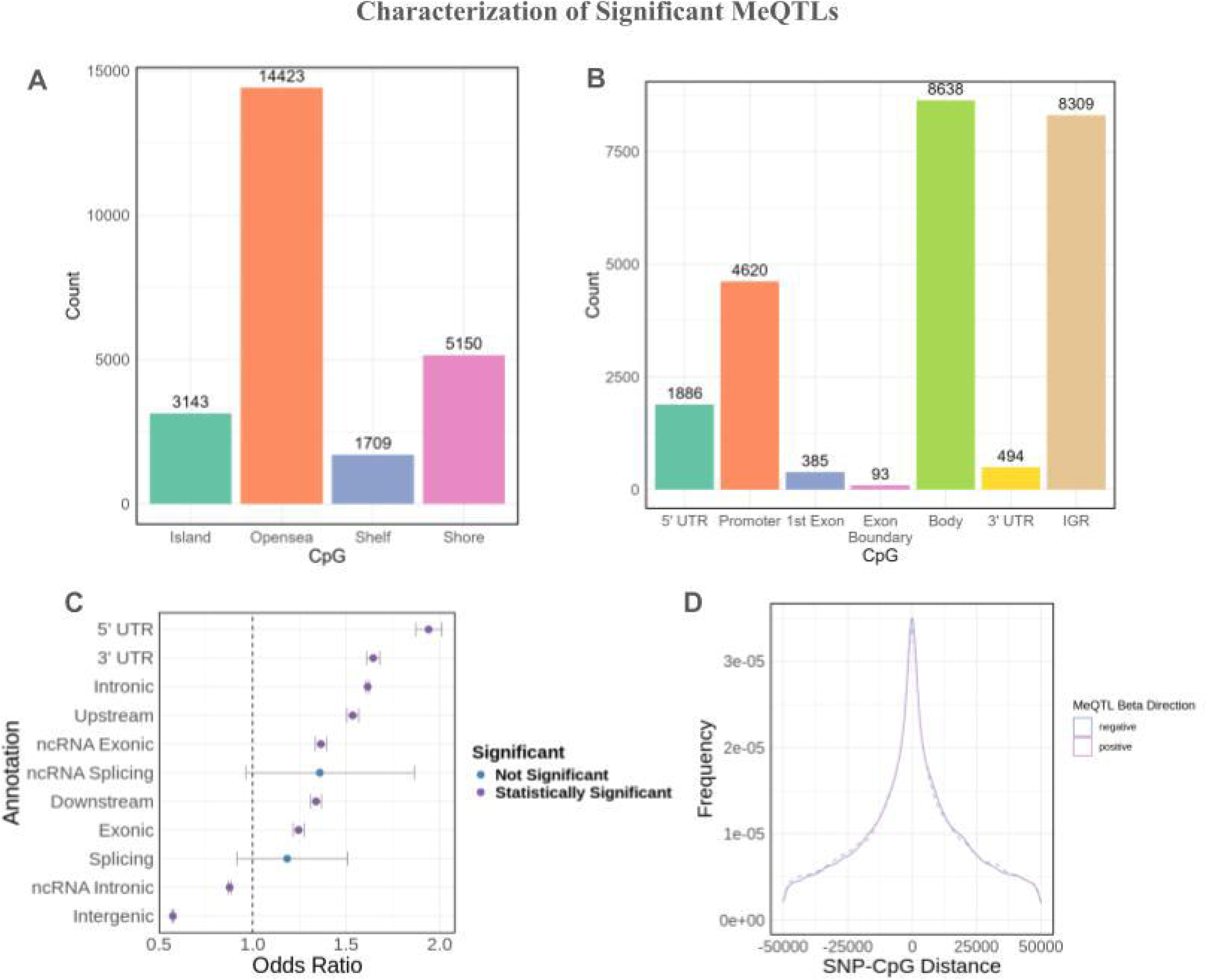
Overview of significant mcQTLs in African American hepatocytes, a) CpG types for significant meQTLs. b) CpG genomic annotations for significant meQTLs. c) Enrichment analysis of SNP annotations for significant meQTLs. d) MeQTL density by SNP-CpG distance.

Given that methylation is a known regulator of gene expression, we were interested in determining if SNPs associated with differential methylation were also associated with differences in gene expression.

### Colocalization Analysis

Many of the variants identified through meQTL mapping are known liver eQTLs. To determine if the same causal variant drives differences in methylation and expression for a given gene, we performed colocalization analysis between our significant meQTLs and eQTL data from two distinct eQTL datasets.

First, colocalization analysis was performed for 61,851 SNPs that were significant meQTLs in our dataset and significant eQTLs in the GTEx V8 liver dataset. Within the GTEx eQTL dataset, these shared SNPs were associated with the regulation of 2,363 eGenes, resulting in a total of 120,874 GTEx eQTLs included in the analysis. At PP4 > 0.8, 18,209 SNPs were colocalized between the two datasets (Figure 2A). In our meQTL dataset, these 18,209 SNPs are associated with differential methylation at 1,559 methylation sites. In GTEx, these SNPs are associated with the expression of 1,122 eGenes (Figure 2B), including 83 genes involved in metabolic pathways and 17 genes involved in drug metabolism and drug resistance. This includes CYP enzymes such as *CYP2C19*, which plays a critical role in converting drugs such as Clopidogrel to their active form,^28^ as well as glutathione transferases such as *GSTM3*, *GSTA4*, *GSTT2*, and *GSTT2B*, which are involved in Phase II drug metabolism and detoxification.^29^ Given that eQTLs for 2,363 eGenes were analyzed in colocalization analysis, 47.48% of the genes tested exhibited colocalization with meQTLs, indicating that methylation may mediate the effects of SNP variation on gene expression for a large proportion of GTEx eGenes. One limitation of this analysis is that GTEx eQTLs are for whole liver tissue and encompass a variety of cell types, while our meQTLs were identified specifically in hepatocytes. Additionally, the GTEx study cohort is primarily composed of European individuals. Due to differences in LD structure and allele frequencies between populations, the GTEx project may fail to capture eQTLs regulating gene expression in non-European individuals.

**Figure 2:**
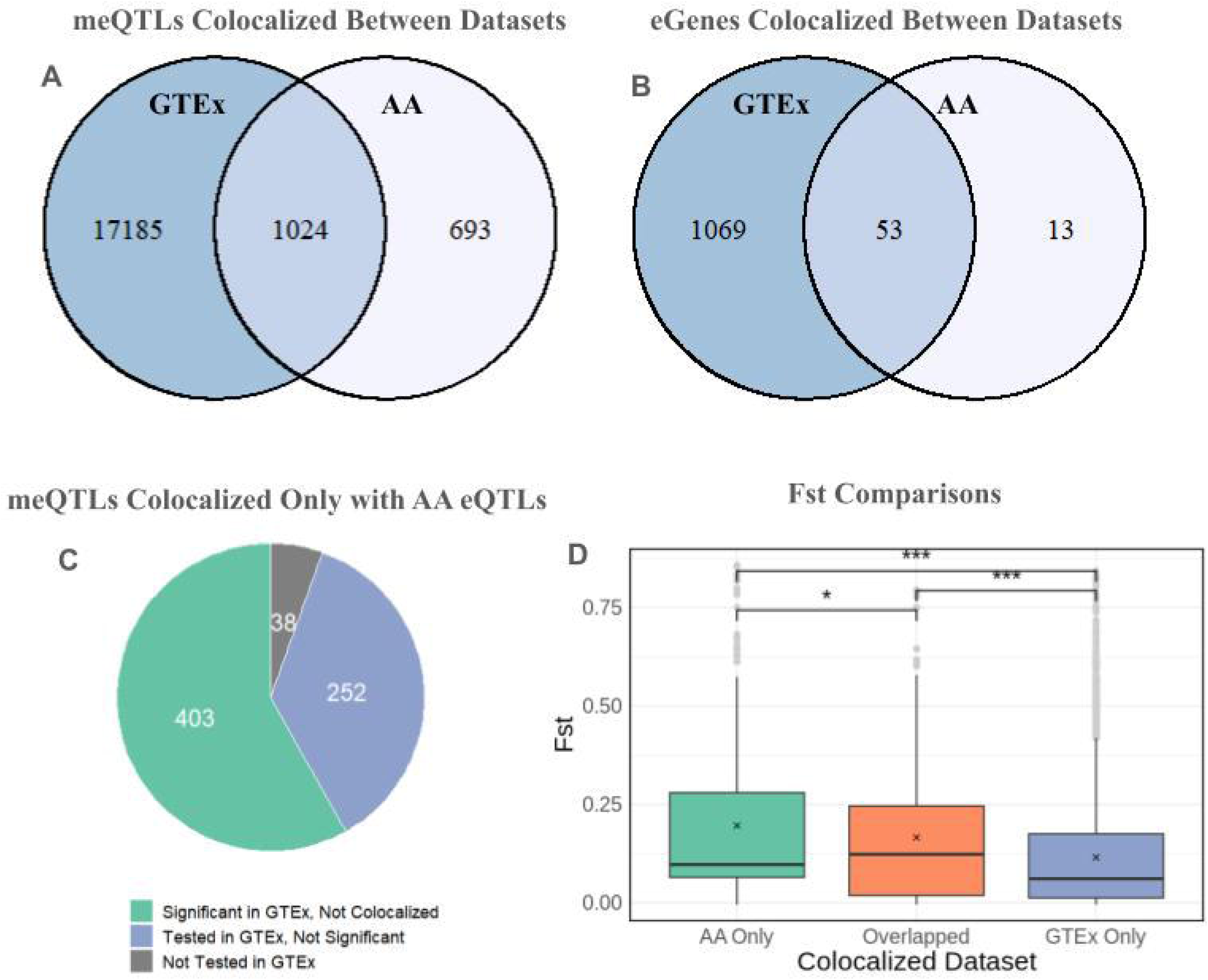
Results from colocalization analysis between African American meQTLs and two eQTL datasets, a) Number of meSNPs colocalized with GTEx eSNPs only (dark blue), AA eSNPs only (white), or both datasets (light blue), b) Number of eGenes that are associated with SNPs only colocalized with GTEx eQTLs (dark blue), AA eQTLs (white), or eQTLs that are colocalized in both datasets (light blue), c) Breakdown of SNPs that arc colocalized with the AA eQTL dataset, but not the GTEx dataset. These SNPs were significant GTEx eQTLs that did not colocalize (green), eQTLs that were tested in GTEx but did not reach statistical significance (blue), or eQTLs that were not tested in GTEx (gray), d) Comparison of Fst values between SNPs that colocalized exclusively with African American eQTLs (green), SNPs that colocalized with both eQTL datasets (orange), and SNPs that colocalized exclusively with GTEx eQTLs (blue).

Because our meQTL data was obtained from an African American cohort, we also performed colocalization with an African American eQTL dataset. 8,614 SNPs exhibit significant associations in both datasets, which corresponds to 188 significant eGenes in the AA eQTL dataset. Of the 8,614 SNPs with significant associations in both datasets, 1,717 of these SNPs exhibited colocalization at PP4 > 0.8 (Figure 2A). These SNPs are associated with differential methylation at 192 CpGs in our meQTL dataset. In our AA eQTL dataset, these colocalized SNPs are associated with the expression of 66 eGenes (Figure 2B). Thus, given that African American eQTLs for 188 eGenes were tested for colocalization, 35.1% of eGenes tested had an eQTL that exhibited colocalization with an AA meQTL. Colocalization with the African American dataset resulted in fewer significant colocalizations compared to the GTEx data. This was expected because the African American eQTL cohort is composed of 60 individuals, thus reducing the number of significant eQTLs identified in this study.

### Comparison of Colocalization Results by Dataset

1,024 SNPs and 53 eGenes exhibited colocalization in both datasets. However, there were notable differences in SNPs and eGenes that were colocalized in the AA eQTL dataset, but not in GTEx. For instance, *F12*, which encodes the coagulation Factor XII protein, is colocalized with the same two SNPs (rs1801020, rs2731674) in both eQTL datasets. These two SNPs are in high LD, with a R^2^ value of 0.8123 in African Americans. Both of these SNPs were associated with methylation at 2 CpGs in the Factor 12 gene body in meQTL analysis. Data for rs1801020 is shown in Figure 3.

**Figure 3:**
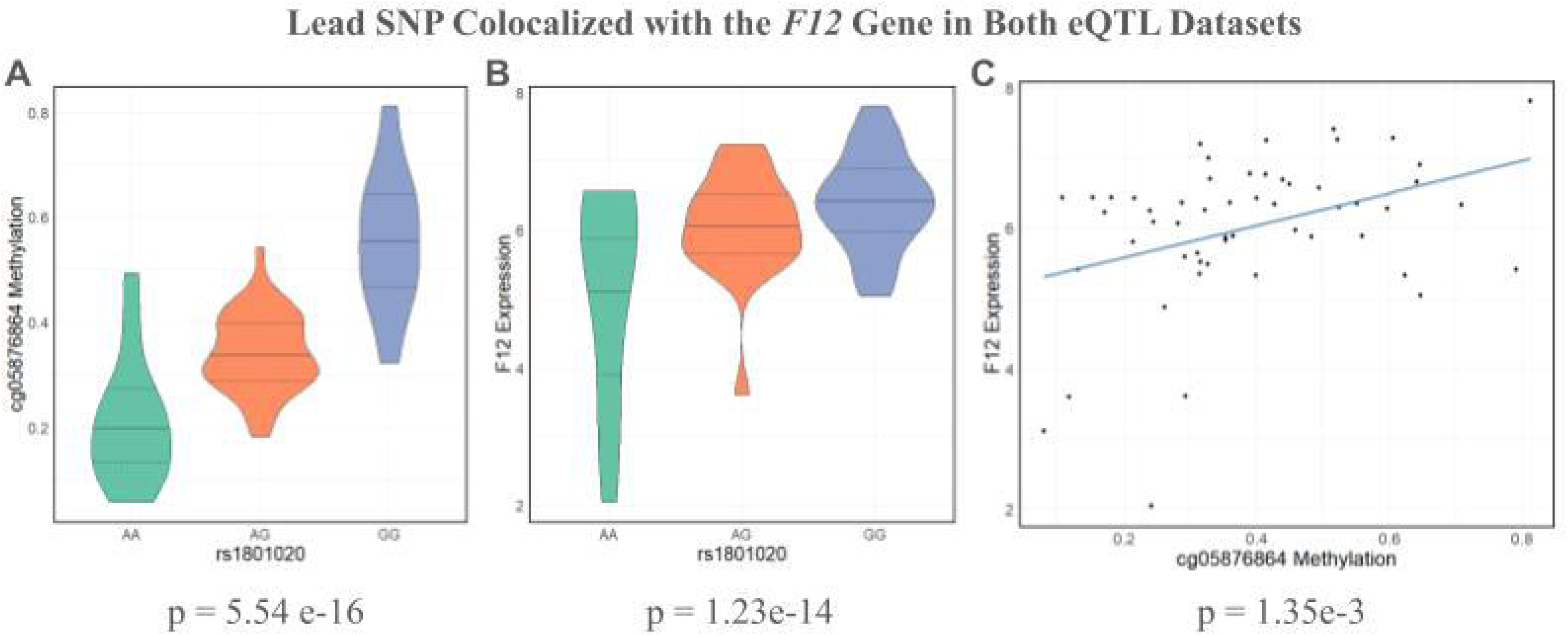
Methylation and expression data from African Americans for rs 1801020. a SNP that colocalizcd with *Factor 12* in both datasets, a) Violin plot of methylation at cgO5876864. the lead CpG site associated with rs 1801020 in meQTL analysis, according to rsl 81020 genotype, b) Violin plot of *F12* expression according to rsl8O!O2O genotype. The G allele is associated with increased methylation and increased gene expression, c) Scatterplot of *F12* expression versus cgO5876864 methylation. Increased methylation at this CpG is associated with increased gene expression.

For both of these SNPs, the G allele is associated with increased methylation and increased gene expression. rs1801020, the lead meQTL for *F12*, had a p-value of 5.54e-16 in meQTL analysis (Figure 3A) and a p-value of 1.23e-14 in eQTL analysis (Figure 3B). rs2731674 had a p-value of 5.26e-13 in meQTL analysis and 2.28e-12 in eQTL analysis. The minor allele for rs1801020 has been linked to decreased thrombotic risk in both African Americans and European Americans.^30^ This SNP lies 4 bases upstream of the transcription start site for *F12*, suggesting that it plays a key role in regulating this gene’s expression.^31^ Though rs2731674 has not been directly implicated in thrombotic risk, genome-wide association studies have identified associations with thrombospondin, Factor IX, and kallikrein levels, all of which are involved in the coagulation response.^32^ This provides strong evidence that Factor XII SNPs may regulate FXII expression and thrombotic risk by altering methylation in this gene.

Conversely, several SNPs and eGenes only exhibit colocalization with African-American eQTLs. 13 eGenes exhibited colocalization with African American eQTLs, but were not colocalized with the GTEx eQTLs. These genes were not strongly enriched for any particular pathway or disease. 1,024 SNPs were colocalized with both GTEx and our African eQTL dataset (Figure 2A). However, there were 693 SNPs that exhibited colocalization with our AA eQTL dataset but *not* with GTEx. As shown in Figure 2D, meQTLs colocalizing only with the AA eQTLs had a greater average fixation index (Fst = 0.196) compared to SNPs colocalizing in both datasets (Fst = 0.165, p = 0.0248) and compared to SNPs colocalizing only in GTEx (Fst = 0.115, p = 3.98 e-18). This suggests that using African American eQTL data for colocalization analysis enables the identification of additional causal SNPs that were not identified in non-African datasets, especially for SNPs that have greater frequency differences between African Americans and their European counterparts.

Of the 693 SNPs that exhibited colocalization with the African-American eQTLs but not with the GTEx eQTLs, 403 of these SNPs were significant GTEx eQTLs (Figure 2C). 252 of these SNPs were tested for association in GTEx eQTL analysis, but did not reach the threshold for statistical significance. As discussed in Zhong et al., it is believed that these eQTLs were only detected in the African American cohort because they exist at higher frequencies in African American populations compared to European populations or are specific to gene regulation in hepatocytes as opposed to liver.^10^ The remaining 38 SNPs were not tested in GTEx, but were significant eQTLs in Zhong et al.^10^

One gene that exhibited striking differences in colocalization between the GTEx and AA eQTL datasets is *HLA-G*, an immune checkpoint molecule that has been implicated in gastrointestinal disease, diabetes, and adverse pregnancy outcomes such as preeclampsia.^31, 32, 33^ 61 meSNPs exhibited colocalization with the GTEx dataset, while 35 meSNPs exhibited colocalization with the AA dataset. Interestingly, there is no overlap in colocalization between the two datasets. As shown in Figure 4A, the eQTL results for *HLA-G* exhibited striking differences between the two eQTL datasets. While the significant eQTLs identified in GTEx are dispersed across a 1.8 Mb region, the significant eQTLs identified in the AA data fall in a narrow 25.6 kB region.

**Figure 4:**
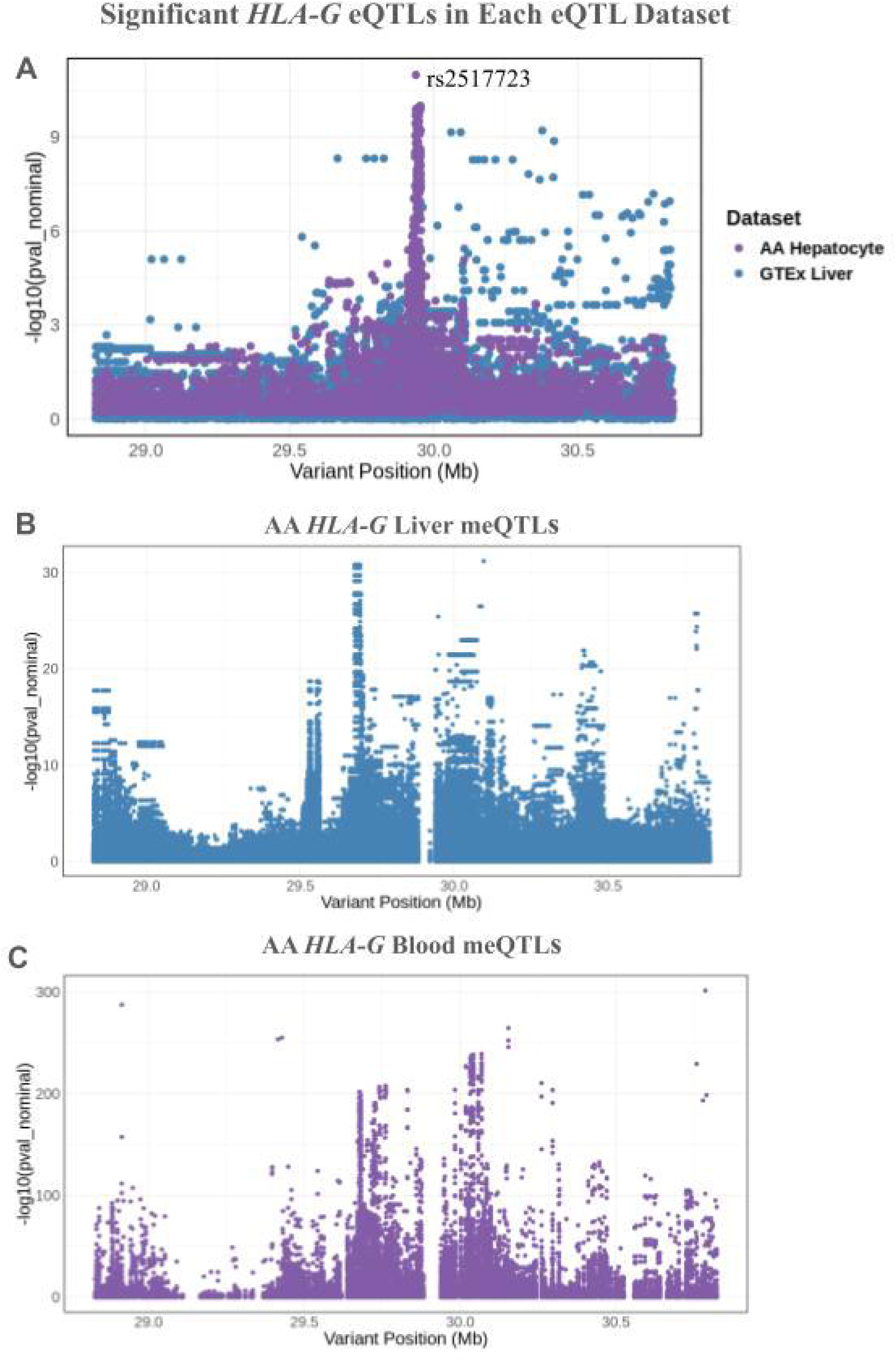
Significant eQTLs for *HLA-G* in the African American and GTEx eQTL datasets, a) Manhattan plot for *HLA-G* eQTLs identified in GTEx liver (blue) and African American hepatocytes (purple). While the GTEx eQTLs are dispersed throughout the 1.8 Mb region, the African American eQTLs are concentrated in a small spike, b) *HLA-G* liver meQTLs identified in African American hepatocytes, c) *HLA-G* blood meQTLs identified in the GENOA study. In Figures 4B and 4C, the Manhattan plots have similar structure, indicating that blood meQTLs for *HLA-G* are largely shared between blood and liver.

Furthermore, the AA eQTLs that exhibit colocalization with meQTLs are located within 15.4 kB of each other, creating a sharp spike in the Manhattan plot. In contrast, the GTEx eQTLs that exhibit colocalization with meQTLs are spread out across a 958 kB region. Due to differences in eQTL structure between populations, the use of African American meQTL data paired with African American eQTL data enhances the identification of causal SNPs regulating DNA methylation and gene expression in this population.

### Direction of Effect Analysis for SNPs Colocalized with African American eQTLs

To better understand the mechanisms through which colocalized meQTL-eQTL SNPs regulate DNA methylation and gene expression, we annotated these SNPs and CpGs to their respective regions in the genome and examined their direction of effect.

Of the 1,717 SNPs colocalized with the African American dataset, 1,337 of these SNPs have discordant meQTL and eQTL betas. This means that the effect allele has opposite directions of effect on DNA methylation and gene expression (ex. the allele is associated with increased methylation and decreased gene expression, or decreased methylation and increased gene expression). These SNPs are associated with CpGs that tend to fall in shore and opensea regions (Figure 5A). Additionally, a high proportion of these CpGs lie in the 5’ UTR or promoter, where methylation can repress the initiation of transcription. A high proportion of CpGs also fall in the gene body (Figure 5B).

**Figure 5:**
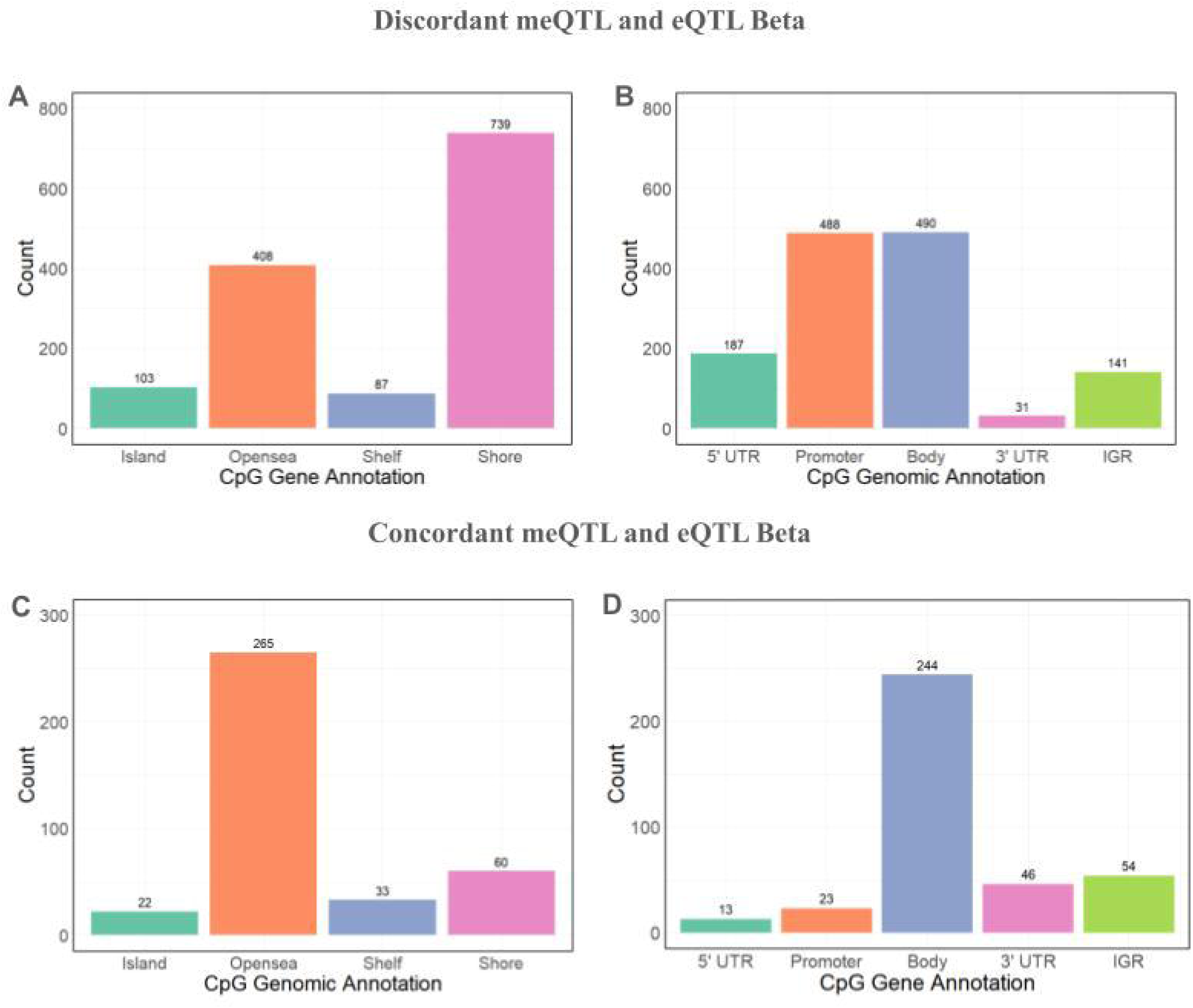
Distributions of CpG annotations for colocalizcd cQTL-mcQTL pairs in African Americans, a) CpG genomic annotations for CpGs where the associated SNP has discordant meQTL and eQTL betas. For these SNPs, a given allele has the opposite direction of effect on methylation and gene expression (ie. increased methylation and decreased expression, or decreased methylation and increased expression). The greatest number of CpGs fall in shore regions, b) CpG gene annotations for colocalizcd CpGs, discordant beta. Most CpGs fall in the gene body, promoter, and 5’ UTR. c) CpG genomic annotations for colocalizcd CpG sites, concordant beta. For these SNPs, a given allele has the same direction of effect on methylation and gene expression (ie. increased methylation and increased gene expression, or decreased methylation and decreased gene expression). Most CpGs fall in opensea regions, d) CpG gene annotations for colocalized CpG sites, concordant beta. Relative to when the meQTL and eQTL betas are discordant, a greater proportion of CpGs fall in gene body and fewer fall in the 5’ UTR and promoter.

The remaining 380 SNPs have concordant meQTL and eQTL betas, meaning the effect allele has the same direction of effect on DNA methylation and gene expression (ex. the allele is associated with both increased methylation and increased gene expression). Compared to when the meQTL and eQTL betas are discordant, a greater proportion of these CpGs fall in the gene body and significantly fewer fall in the promoter (Figure 5D). For instance, for the *Factor 12* gene, which encodes Coagulation Factor XII and is involved in anticoagulant drug response, increased methylation in the gene body is associated with increased gene expression (Figure 3C). While methylation is often viewed as a repressive transcriptional marker, especially when deposited near the promoter, this analysis demonstrates that methylation, particularly in the gene body, can also be positively associated with increased gene expression.

### Mediation Analysis

To determine the extent to which methylation mediates the effect of SNP genotype on gene expression, we performed mediation analysis for 1,122 CpG-gene pairs that were colocalized at PP4 > 0.80. We performed mediation analysis using two models. First, we assessed the extent to which methylation mediates the effect of SNP genotype on gene expression (SNP-Methylation-Expression analysis or “SME”). Second, we assessed whether gene expression mediates the effect of SNP genotype on DNA methylation (SNP-Expression-Methylation analysis or “SEM”). At a FDR of 0.05, we identified a total of 34 CpG-gene pairs with significant evidence for mediation in either of these models. Of these 34 pairs, 29 CpG-gene pairs were significant in both analyses, 1 was significant in SME analysis but not SEM, and 4 were significant in SEM analysis but not SME. At a FDR of 0.10, we identified 63 CpG-gene pairs with significant evidence for mediation. 53 CpG-gene pairs were significant in both analyses, 5 were significant in SME analysis but not SEM, and 5 were significant in SEM analysis but not SME.

We identified two meCpG-eGene pairs of interest that are associated with Type 1 Diabetes (T1D), a disease whose prevalence is increasing in ethnic minorities such as African Americans.^36^ The first meCpG, cg00030291, was found to mediate the effect of rs3779356 on the expression *ICA1*, a gene whose hypermethylation and differential expression have been previously linked to diabetes.^37^ This SNP was colocalized only with the GTEx dataset, as this was not a significant eQTL in the AA eQTL dataset. The second meCpG, cg13872627, likely mediates the effect of rs7767978 on *HLA-C* expression, another gene that has been linked to T1D.^38^ rs7767978 was colocalized with the African American eQTL dataset. Despite being a significant eQTL for HLA-C in GTEx, this SNP was not colocalized with the GTEx eQTL dataset at a statistically significant level.

### GWAS Enrichment Analysis for SNPs Colocalized with AA eQTLs

Given that nearly 90% of variants identified in previous genome-wide association studies (GWAS) lie in non-coding regions of the genome and may regulate gene expression through epigenetic modifications, we were curious to see whether meSNPs colocalized with the African American eQTL dataset are also SNPs that have been associated with disease in previous GWAS. Additionally, we were interested in determining if any particular parent Experimental Factor Ontology (EFO) categories were enriched in our analysis, as this would indicate that the SNPs identified through colocalized analysis may contribute to these diseases by affecting DNA methylation.

Our colocalized SNPs overlapped with GWAS hits for 9 parent EFO categories. After performing EFO category enrichment, we identified 5 parent EFO terms that are enriched for colocalized meQTL-eQTL SNPs at statistically significant levels and 1 parent EFO term that is significantly depleted in our list of colocalized SNPs (Figure 6) . We observed an enrichment for inflammatory measurements (OR = 4.509), lipid or lipoprotein measurements (OR = 4.184), and digestive system disorders (OR = 2.096), suggesting that methylation plays a role in regulating genes associated with these diseases.

**Figure 6:**
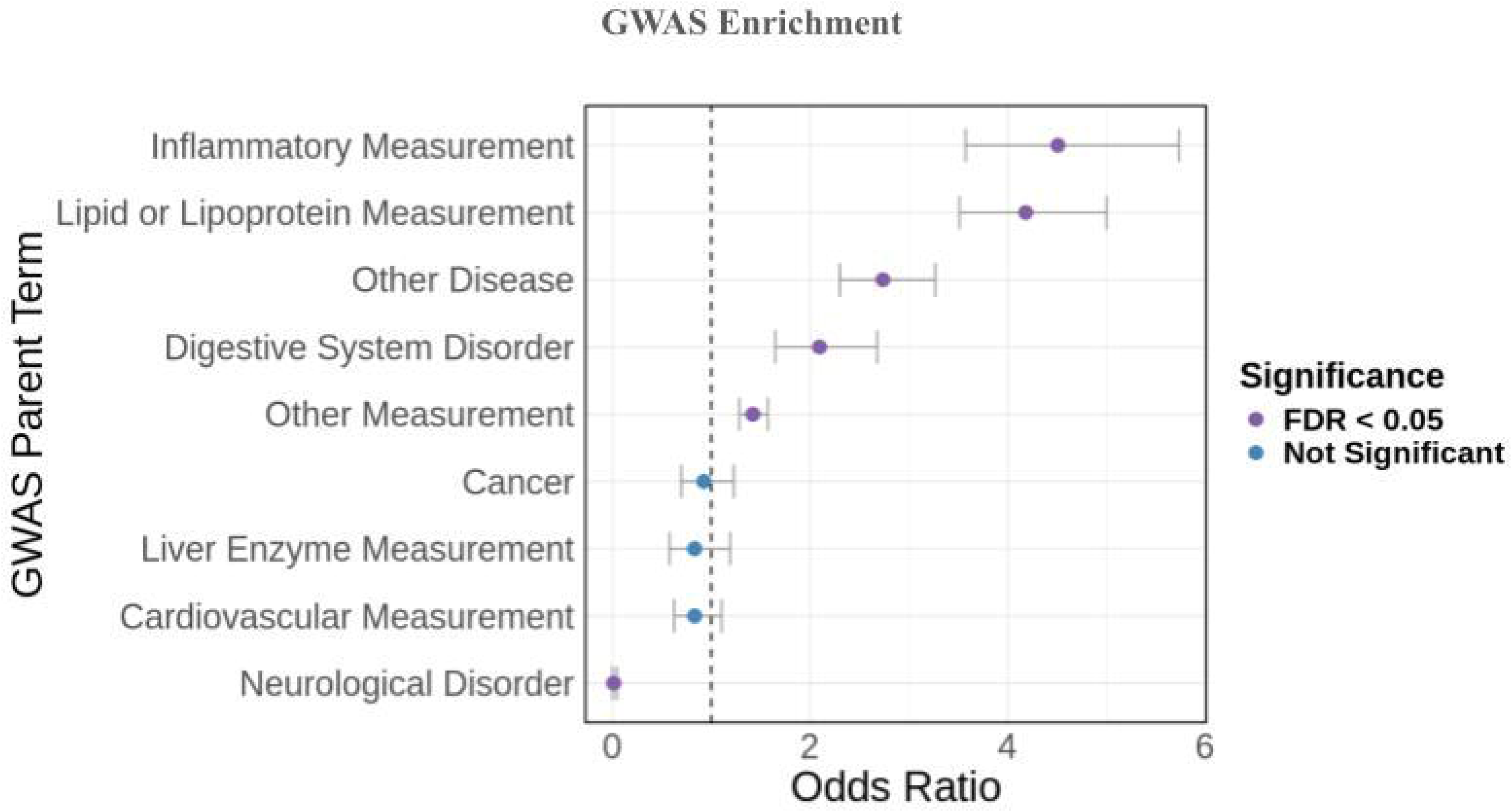
GWAS Parent EFO term enrichment for meSNPs colocalizcd with the African American eQTL dataset. Colocalized SNPs were associated with 9 different parent EFO categories, and 6 of these categories exhibited enrichment or depiction at statistically significant levels. Parent EFO categories arc ordered by odds ratio. Error bars represent the 95% confidence interval from Fisher’s test.

### Colocalization with AA MeQTLs in Blood

Finally, to determine whether the same variants regulate CpG methylation and gene expression in both liver and blood, we performed colocalization analysis with our African American liver meQTLs and African American blood meQTLs identified in the GENOA study.^9^ Several studies have used methylation in the blood as a biomarker for various liver diseases. However, the presumption of shared methylation effect is not well-tested in African Americans.^39, 40, 41^ If the same variants regulate DNA methylation in both liver and blood, this would suggest that biomarkers informative of gene expression in blood are also robust indicators of gene expression in the liver. As discussed previously, differences in the ancestral makeup of study cohorts can affect colocalization results. Because our meQTL study and the GENOA meQTL study both investigated African American populations, any observed differences in colocalization between AA blood and liver would suggest that the regulation of DNA methylation and gene expression is distinct to each tissue.

The GENOA meQTL project identified a total of 4,565,687 significant blood meQTLs, while we identified 410,186 significant liver meQTLs. Of the liver meQTLs we identified, 264,797 SNPs are also significant AA blood meQTLs. However, colocalization analysis revealed that only 14,496 SNPs, or 5.4% of all SNPs tested, are colocalized between the two datasets at PP4 > 0.80. Some of these colocalized SNPs fall in genes involved in the absorption, detoxification, metabolism, and excretion (ADME) of drugs. For instance, one meQTL (rs276204) in *GSTO2*, a glutathione S-transferase involved in the detoxification of xenobiotics,^29^ exhibits colocalization between liver and blood. Additionally, 4 meQTLs (rs3094159, rs3132718, rs12527415, rs141143417) regulating methylation near the *HLA* genes exhibit colocalization in blood and liver, suggesting that the regulation of methylation in these genes is shared between blood and liver. Figures 3B and 3C further demonstrate that the Manhattan plots for *HLA-G* meQTLs are similar between blood and liver, suggesting that the same SNPs regulate this gene’s methylation in both tissues. For the 250,301 liver meSNPs that did not exhibit colocalization with blood meQTLs, knowing these variants’ effects on DNA methylation in blood cannot inform our understanding of the genetic regulation of methylation in the liver.

## Discussion

In this study, we performed cis-meQTL mapping in African American hepatocytes to identify cis-SNPs regulating DNA methylation in these liver cells. By integrating this analysis with eQTL data from a predominantly European cohort (GTEx) and an African American cohort, we identified a total 18,209 colocalized SNPs that regulate both DNA methylation and gene expression. The SNPs colocalized with African American eQTL data exhibit a significant enrichment for GWAS EFO terms such as lipid or lipoprotein measurement, inflammatory measurements, and gastrointestinal disorders, indicating that these SNPs, as well as the methylation sites they affect, may be valuable biomarkers for these diseases.

Most importantly, we demonstrated that performing epigenomic studies in an African American population can reveal novel genetic regulators of gene expression. While nearly 60% of meSNPs colocalized with the African American eQTL dataset were also colocalized with the GTEx dataset, we identified 693 variants that are only colocalized with the AA eQTLs. Due to differences in LD between African and European American populations, using African American eQTL data allows us to identify a population-specific set of potentially causal variants. Our analysis of *HLA-G* meQTL colocalization with GTEx showed that eQTL associations for this gene were dispersed throughout a 1.8 Mb region, while eQTL associations for this gene in the African American eQTL dataset were concentrated in a narrow spike spanning a 25.6 kB region (Figure 4A). A similar association pattern was observed when comparing significant meQTL associations between blood and liver in African Americans (Figure 4B, 4C). In both blood and liver, the meQTL associations tend to be clustered within a narrow genomic region rather than spread out across the gene. Furthermore, the pattern of these associations in the liver meQTL Manhattan plot is strikingly similar to the pattern in the blood meQTL plot. This narrow spiking pattern was replicated across the African American liver eQTL data, liver meQTL data, and blood meQTL data but not in the GTEx eQTL data, suggesting this effect is driven by the unique genetic architecture of African Americans.

Furthermore, African American meQTLs that exhibit colocalization with African American eQTLs are enriched for clinically relevant GWAS EFO terms, such as lipid or lipoprotein measurement. By limiting our analysis to colocalized variants, we are adding potential mechanistic insight to how these significant GWAS variants regulate gene expression via DNA methylation. African Americans face an increased risk of developing cardiovascular disease, as well as an increased mortality rate compared to their white counterparts.^42^ The liver is critical in the homeostasis of plasma lipids, as it responds to fasting and feeding by synthesizing, storing or secreting lipids. The genes regulating these processes can play an important role in disease susceptibility and progression. Thus, identifying population-specific genetic drivers of cardiovascular disease is critical for understanding why African Americans exhibit an increased risk of developing these conditions and could guide future investigations of African American-specific genetic risk factors for these diseases.

Through colocalization analysis between blood and liver meQTLs, we demonstrated that for some genes, the SNPs regulating methylation in blood also regulate methylation in the liver. For instance, the same SNPs regulate *GSTO2* methylation in African American blood and livers. In a previous study investigating the genetic and epigenetic regulation of gene expression in livers, it was found that methylation at cg23659134, the same methylation site that exhibited colocalization in our analysis, explains a greater proportion of variability in *GSTO2* expression than SNP genotypes alone.^43^ Our analysis further adds to the existing knowledge base of methylation patterns in genes involved in drug metabolism by demonstrating that the same *GSTO2* SNPs regulate methylation at this CpG site in both blood and liver. Furthermore, our results would indicate that studying *GSTO2* methylation in blood can provide additional insight into this gene’s regulation in the liver. Using blood biomarkers to predict *GSTO2* activity in the liver could potentially streamline studies of drug metabolism by eliminating the need to assay liver-specific gene expression.

This study has several limitations. First, our meQTL study cohort was relatively small, with only 77 hepatocyte donors participating in this study. This likely decreased our power to detect relevant hepatocyte meQTLs. In addition, the AA liver eQTL dataset we used for colocalization analysis also had a relatively small sample size, with only 60 donors, which reduced the number of eQTLs that could be tested for colocalization. This explains why colocalization analysis with the GTEx dataset identified thousands of colocalized meQTL-eQTL variants, while colocalization with the AA dataset identified fewer than 2,000 colocalized variants. Despite the limitations of small cohort sizes, this study still represents a major step in improving our understanding of gene regulation in African Americans and increasing the availability of epigenomic data from African American populations.

To continue developing our understanding of how genetic variants regulate DNA methylation and gene expression in African Americans, several lines of future research are possible. Massively parallel reporter assays are a valuable tool for efficiently interrogating the regulatory effects of candidate variants *in vitro* and could be used to functionally validate the candidate variants identified in this study. Further investigating how genetic factors regulate DNA methylation and gene expression in African Americans will provide additional insight into the molecular mechanisms underlying disease risk and drug response in this group and improve health outcomes for this understudied population.

## Supporting information

Colocalization Analysis Summary Statistics, African American eQTLs

Colocalization Analysis Summary Statistics, GTEx eQTLs

## Author Contributions

Writing, data curation, and formal analysis were conducted by KC. Data curation and methodology were conducted by KC, CC, YZ, GY, CA and MM. Conceptualization, funding acquisition, supervision, and writing were conducted by MAP.

## Data Availability Statement

Results from colocalization analysis are available under Supplementary Data. Code generated during this study is available at https://github.com/pereralab. Summary statistics from cis-meQTL analysis will be made available on FigShare.

---

**Supplementary Figure 1:**
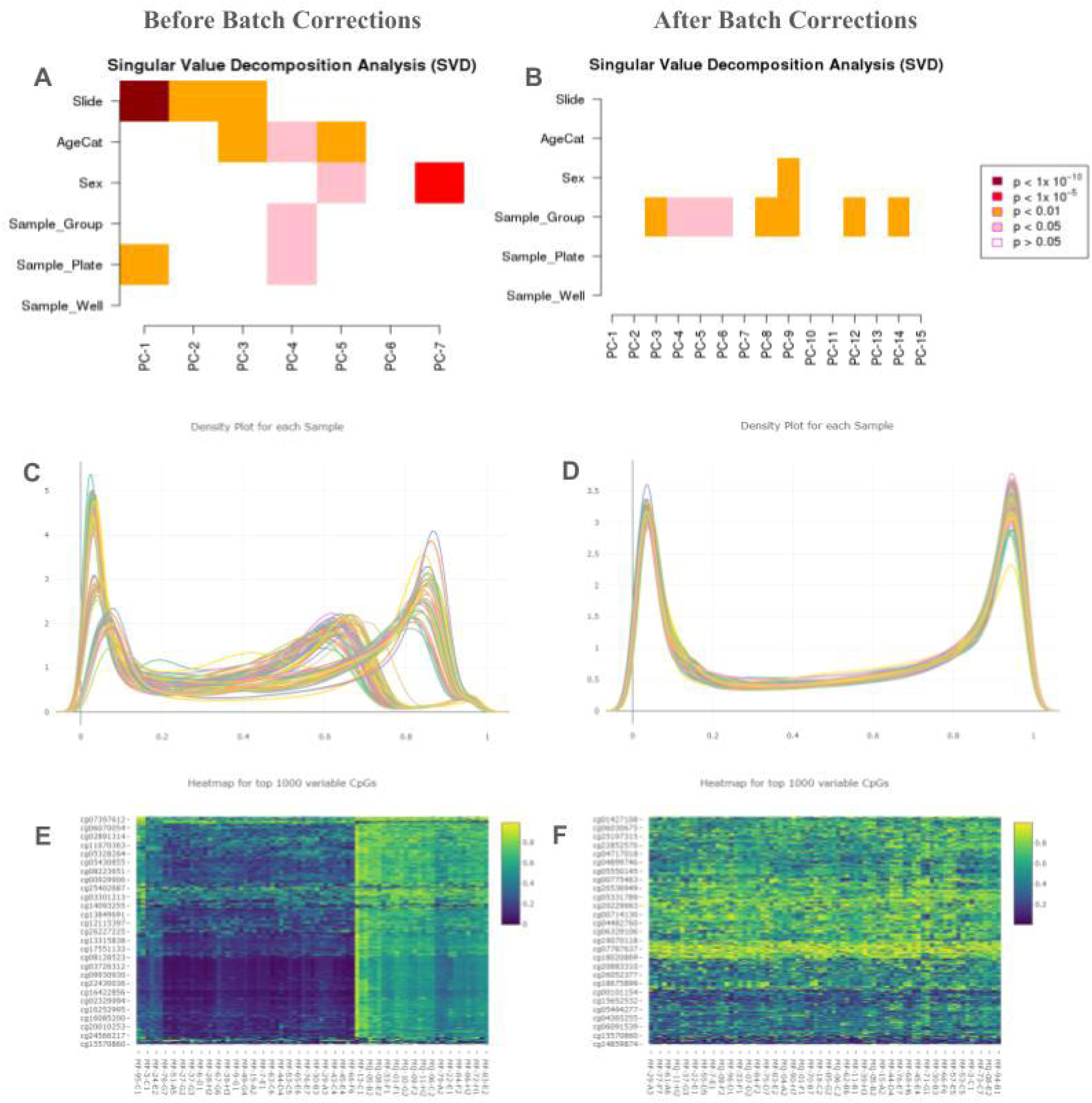
Reduction of batch effects using the *ChAMP* package in R. a, b) Use of the ChAMP.runCombat function reduces the effects of technical variation, such as slide and sample plate, on methylation measurements, c, d) Prior to normalization, methylation measurements fall in three clusters on the density plot. Each cluster represents a separate sequencing batch. Following normalization and corrections for batch effects, the distributions for each batch align, e, f) Prior to normalization, variability in methylation at the top CpGs is primarily driven by batch effects, as there are strong differences between the clusters of samples. Following normalization. all samples display similar variability at these CpGs. These three figures demonstrate that *ChAMP* sufficiently corrected for batch effects to reduce the effects of technical variation on methylation measurements.

